# The Rho GTPase exchange factor Vav2 promotes extensive age-dependent rewiring of the skin stem cell transcriptome

**DOI:** 10.1101/2022.09.27.509634

**Authors:** L. Francisco Lorenzo-Martín, Xosé R. Bustelo

## Abstract

Both the number and regenerative activity of skin stem cells (SSCs) are regulated by Vav2, a GDP/GTP exchange factor involved in the catalytic stimulation of the GTPases Rac1 and RhoA. However, whether Vav2 signaling changes in SSCs over the mouse lifespan is not yet known. Using a mouse knock-in mouse model, we now show that the expression of a catalytically-active version of Vav2 (Vav2^Onc^) promotes an extensive rewiring of the overall transcriptome of SSCs, the generation of new transcription factor hubs, and the synchronization of many transcriptional programs associated with specific SSC states and well-defined signaling pathways. Interestingly, this transcriptome rewiring is not fixed in time, as it involves the induction of 15 gene expression waves with diverse distribution patterns during the life of the animals. These expression waves are consistent with the promotion by Vav2^Onc^ of several functional SSC states that differ from those normally observed in wild-type SSCs. These results further underscore the role of Vav2 in the regulation of the functional state of SSCs. They also indicate that, unlike other Vav2-dependent biological processes, the signaling output of this exchange factor is highly contingent on age-dependent intrinsic and/or extrinsic SSC factors that shape the final biological readouts triggered in this cell type.

**AUTHOR SUMMARY:** Skin stem cells (SSCs) are essential for the homeostatic balance of the skin, yet little is known to date about the biological and molecular mechanisms that modulate their abundance, long-term stability, or functional states during ageing. To address this issue, in this work we have used a genetically-engineered gain-of-function mouse model for Vav2, a Rho guanine nucleotide exchange factor (GEF) that has been recently shown to be involved in skin stem cell homeostasis. By performing time-course genome-wide expression analyses combined with a number of computational methods, here we show that: (i) Vav2 plays a critical role in regulating the functional state of SSCs, and (ii) the signaling output of constitutively active Vav2 is highly contingent on age-dependent intrinsic and/or extrinsic SSC factors that shape the final biological readouts triggered in this cell type. We believe that these data represent, to our knowledge, one of the first examples of the time-dependent output of an oncogenic version of a Rho GEF along a wide time interval in mice.

## INTRODUCTION

SSCs are essential for the homeostatic balance of the skin (1, 2). They are located in the bulge area of the hair follicle, from which they migrate and progressively differentiate to give rise to the interfollicular, sebaceous, and hair follicle lineages when the skin has to be regenerated due to wounds or other traumas (1-4). Consequently, alterations in the number and/or normal functions of these cells contribute to skin regenerative defects, tumorigenic processes, and ageing (5-7). It is therefore of paramount importance to decipher the biological and molecular mechanisms that modulate their numbers, long-term stability, and functional states.

Most Rho GTPases work as molecular switches depending on the type of guanosine nucleotide bound to them (8). When bound to GDP, these proteins are in an inactive conformation that is not compatible with proximal downstream effector binding. In addition, Rho GTPases are sequestered in the cytosol due to their physical interaction with Rho GDP dissociation inhibitors. However, when in the GTP-bound state, Rho GTPases are unleashed from Rho GDP dissociation inhibitors, interact with a large spectrum of proximal effectors, and eventually promote the stimulation of a wide variety of intracellular functions, such as cytoskeletal remodeling, proliferation, and differentiation (8). The transition of Rho GTPases from the GDP-bound to the GTP-bound state is catalyzed by GDP/GTP exchange factors (GEF), while the reverse inactivation step is mediated by the GTPase activating proteins (8). Numerous Rho GTPase have been implicated in regulation of SSC functions; for instance, Rac1 plays roles in the maintenance of a fully functional SSC reservoir (9-13), RhoA mediates the proliferation and migration of SSCs (14, 15), and Cdc42 regulates the fate of epidermal progenitor cells (16). More recently, we demonstrated that Vav2, a GEF for Rac1 and RhoA, contributes to regulate the number, responsiveness to stimuli, and functionality of SSCs (17). This regulation is associated with transcriptional programs that impinge on the proliferation, pluripotency, and quiescence of SSCs in early postnatal periods (e.g., in 2.5-month-old mice) (17). Consistent with this, we found that the expression of a constitutively active version of Vav2 (referred to hereafter as Vav2^Onc^) in a knock-in mouse model leads to increased numbers of SSCs and more potent skin regenerative responses upon wound healing or hair depilation (17). The opposite effects are observed in mice lacking Vav2 or the related Vav3 protein. Vav2^Onc^ also changes the transcriptome program of the skin cancer stem cells, including the activation of gene signatures directly associated with the undifferentiated and quiescent state of SSCs (17).

We have previously shown that the transcriptional program of SSCs changes quite extensively from early postnatal phases to the adult period in mice (18). Specifically, we have detected several age-specific gene expression waves associated with functions related to proliferation, differentiation, and aging (18). These results raise several intertwined questions regarding the impact of Vav2^Onc^ on the SSC transcriptome: does Vav2^Onc^ promote an age-independent transcriptional program based on its chronically activated state? Alternatively, does Vav2^Onc^ drive a transcriptional program that is subject to age-dependent changes, analogous to those found in normal SSCs (18)? If so, does Vav2^Onc^ only promote the amplification of the normal gene expression waves in SSCs, or does it completely rewire them? To tackle these issues, we have now performed genome-wide expression analyses on SSCs purified from control or Vav2^Onc^-expressing mice at six different time points of aging (ranging from 18 days to 12 months). By coupling this approach to the use of several *in silico* techniques, we obtained an in-depth overview of the age-dependent dynamics of the transcriptome of SSCs in both genotypes.

## RESULTS

### Vav2^Onc^ regulates stem cell transcriptome dynamics

To investigate the effect of the catalytic hyperactivation of Vav2 in SSCs, we decided to monitor the time-dependent evolution of the transcriptome of SSCs isolated from wild-type (WT) and *Vav2*^Onc/Onc^ knock-in mice. The latter mouse strain, which has been described in previous publications from our lab (19, 20), expresses a Vav2 mutant protein (herein, Vav2^Onc^) that exhibits constitutive GEF activity due to the removal of the N-terminal calponin-homology (CH) and acidic (Ac) domains (**Fig. 1A**). These two regions play critical roles in establishing the intramolecular interactions that inhibit the catalytic and most adaptor functions of Vav2 when the protein is not tyrosine-phosphorylated (21-24). Importantly, the mutant allele that encodes Vav2^Onc^ is under the control of the endogenous *Vav2* promoter, thus ensuring that the mutant protein has a tissue distribution and expression levels comparable to those found in the case of the WT counterpart (19, 20). As Vav2^onc^ maintains the adaptor functions that are mediated by the C-terminal SH3–SH2–SH3 cassette of the protein (**Fig. 1A**), the effects elicited by its expression must be catalytic-dependent. Consistent with this, we have previously shown that *Vav2*^Onc/Onc^ mice exhibit phenotypes opposite to those found in mice expressing a catalytically impaired Vav2 protein (L332A mutant) (25, 26).

**FIGURE 1.**
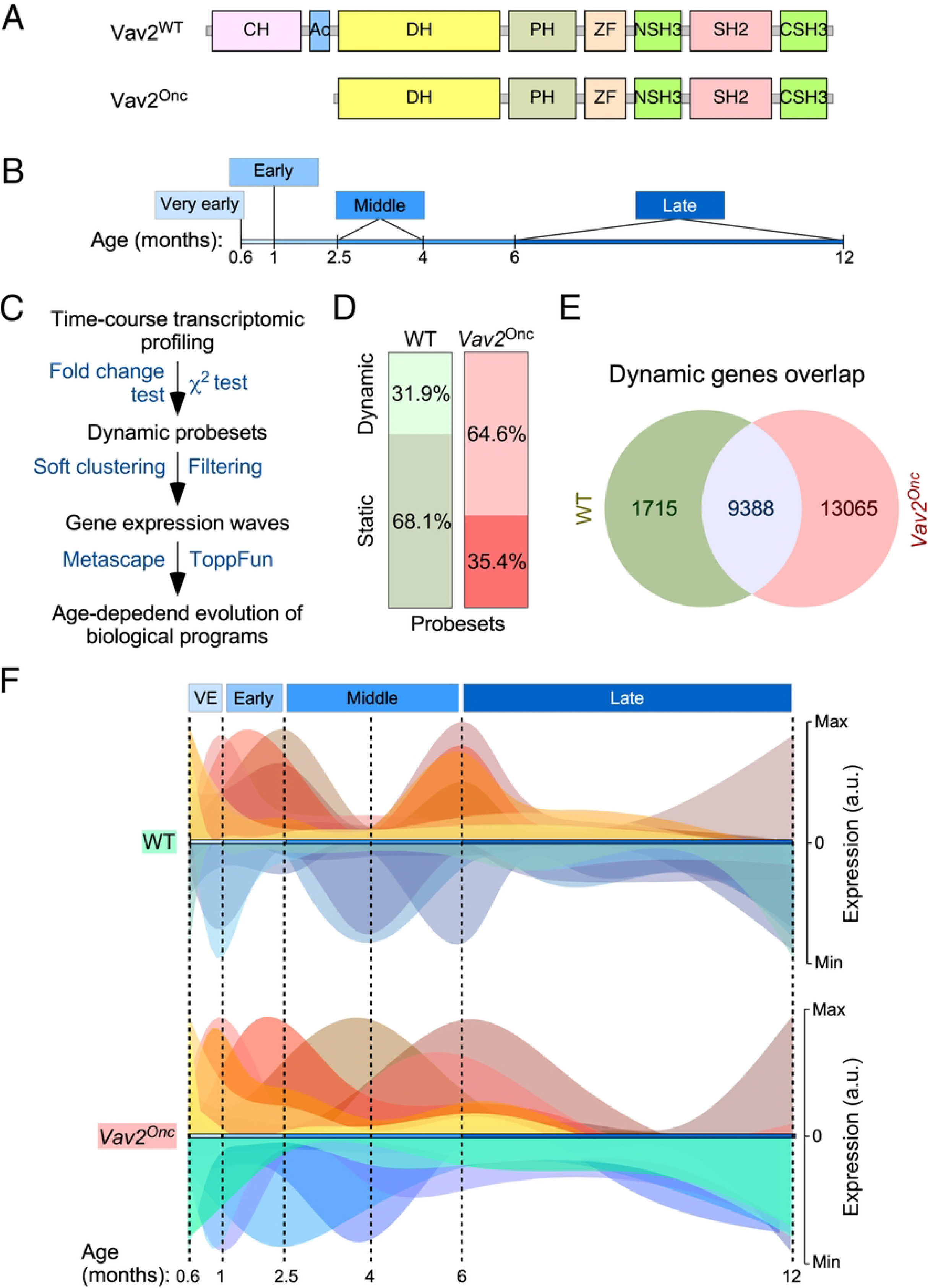
Vav2^Onc^ regulates SSC transcriptome dynamics. **(A)** Scheme of the structure of full-length Vav2 and Vav2^Onc^. Ac, acidic; CH, calponin-homology; CSH3, C-terminal Src-homology 3; DH, Dbl-homology (catalytic domain); NSH3, most N-terminal Src-homology 3; PH, pleckstrin-homology; SH2, Src-homology 2; WT, wild-type ZF, zinc finger of the C1 subtype. **(B)** Time-windows selected to carry out the microarray analysis using samples from SSCs. **(C)** Wet-lab and *in silico* pipeline used to identify the time-dependent transcriptome of SSCs. **(D)** Distribution of the probe sets of the Affymetrix Mouse Gene 1.0 ST array between the dynamic and static groups of SCCs of indicated genotypes. **(E)** Venn diagram showing the overlap in the dynamic probe sets found in the transcriptome of WT and *Vav2*^Onc/Onc^ SSCs. **(F)** Graphical representation of the gene expression waves found in WT (top panel) and *Vav2*^Onc/Onc^ (bottom panel) SSCs. The diagram for the gene expression waves of WT SSCs was taken from (18).

We used flow cytometry to isolate SSCs (CD34^+^ Itgα6^+^) from mice at six time points at different ages (given in months, M), with one time point each classified as very early (0.6-M; 18 days) or early (1-M), and two time points each classified as middle (2.5-M and 4-M) or late (6-M and 12-M) (**Fig. 1B**). The 0.6-M time point corresponds to the earliest stage at which the CD34^+^ Itgα6^+^ population can be clearly identified by flow cytometry (18). Importantly, we did not find any differences in hair cycle staging between WT and *Vav2*^Onc/Onc^ mice (data not shown). Genome-wide expression analyses of total RNA isolated from SSCs were followed by computational approaches specifically designed to identify genes undergoing statistically significant expression changes in at least one of the six time points interrogated (**Fig. 1C**). As previously described (18), these analyses showed that 31.9% of all the mRNAs expressed in WT SSCs display a dynamic behavior across time, according to both *χ*^2^ distribution and expression fold-change (≥ 2-fold) criteria (**Fig. 1D**, left column) (18). In contrast, *Vav2*^Onc/Onc^ SSCs exhibited a much higher percentage (64.6%) of dynamically regulated genes according to the same selection criteria (**Fig. 1D**, right column). These include 85% of the dynamic transcriptome of WT SSCs plus an additional subset of 13 065 probe sets that are specific for *Vav2*^Onc/Onc^ SSCs (**Fig. 1E**).

We next performed soft clustering to visualize all the dynamically regulated genes that were grouped together in coherent gene expression waves (**Fig. 1F**). This was coupled with the use of additional *in silico* tools to allow the identification of the biological processes (GO terms) that were associated with each of those gene expression waves. We found that the SSCs from WT (14 waves) and *Vav2*^Onc/Onc^ mice (15 waves) exhibit similar numbers of time-dependent gene expression waves (**Fig. 1F, Table S1**). These waves are deconvoluted in **Figure 2** to better display the characteristics of each. In this figure, we present the waves found in WT (left columns, dark green) and *Vav2*^Onc/Onc^ SSCs (right columns, dark red) that are classified according to the main time point at which the wave reaches the maximal expression peak value. Each wave is designated with a + or a – symbol depending on whether it contains upregulated (light red waves) or downregulated (light blue waves) genes. In addition, each wave was assigned a reference number that indicates the time point of the maximal expression peak of the wave (e.g., [+6] indicates maximal upregulation of gene expression at 6 months). The reference numbers for multi-peak waves include the number of each peak, always placing the highest fold-change peak in the first position) (e.g., [+1+2.5+6]). Finally, we have included the total number of genes and main GO terms associated with each identified wave (with the size of the letters directly related with the *P* value of each identified GO term) (**Fig. 2**). We found that most of the identified waves in both genotypes have more than one gene expression peak (e.g., waves [+0.6+6], [+2.5+6], [+1+2.3+6], [–1–4], or [–4–12] found in WT SSCs) (**Fig. 2**). The only exceptions to this trend are waves that distribute within the very early time points (e.g., waves [+0.6] and [+0.6+1] in both genotypes) and the late time point (wave [–12] in both genotypes) (**Fig. 2**). However, we also found that the waves in WT SSCs or *Vav2*^Onc/Onc^ SSCs are quite different in terms of size (number of genes involved), shape (distribution in time), and encoded functions (**Fig. 2**). For instance, we observed that the waves in WT SSCs contain similar gene content in the time-windows analyzed (very early phase: 3 763 probe sets [34.9%]; early phase: 2 455 probe sets [22%]; middle phase: 1 162 probe sets [10.8%]; late phase: 3 405 probe sets [31.6%]) (**Fig. 2**). In contrast, in the *Vav2*^Onc/Onc^ SSCs, about half of all the dynamically regulated genes are concentrated in the “late” time point (13 425 probe sets, 59.6%), while the rest of phases exhibit much lower gene content (very early: 4 061 probe sets [18%]; early: 2 485 probe sets [11%]; middle: 2 542 probe sets [11.3%]) (**Fig. 2**). The distribution of the up- and downregulated genes also changes depending on the genotype of the analyzed SSCs. Accordingly, the ratio of upregulated *versus* downregulated probe sets at the late time point is much higher in *Vav2*^Onc/Onc^ SSCs (9 913 vs 3 512, 2.8 ratio) than in WT SSCs (1 489 vs 1 916, 0.8 ratio). Inversions in the ratios of up- and downregulated genes are also seen at the very early (1.2 vs 0.62 ratios) and early (0.82 vs 1.3) phases between *Vav2*^Onc/Onc^ SSCs and WT SSCs (**Fig. 2**). In contrast, the ratios for genes that are mainly regulated in the middle phase are more similar (0.8 and 0.7, for *Vav2*^Onc/Onc^ SSCs and WT SSCs, respectively). Overall, the gene expression waves induced in Vav2^Onc^ SSCs are more enriched in upregulated genes than those in WT SSCs. Finally, we have observed that the wave shape differs greatly between WT and *Vav2*^Onc/Onc^ SSCs (**Fig. 2**). This feature is most conspicuous for the waves of upregulated gene expression at the middle and late time points, which mostly follow a bimodal or a single-peak pattern for WT SSCs or *Vav2*^Onc/Onc^ SSCs, respectively (**Fig. 2**). These shape changes can be illustrated by the single peak–enriched waves of *Vav2*^Onc/Onc^ SSCs at [+2.5], [+4], and [+6], which are in sharp contrast to the frequently observed bimodal waves of WT SSCs at [+1+2.5+6], [+2.5+6], and [+6+2.5] (**Fig. 2**). Interestingly, this disparity in wave shape is less accentuated at the very early and the late time points (see, for example, the waves [+0.6], [+0.6+1], and [–12] from WT or *Vav2*^Onc/Onc^ SSCs; **Fig. 2**). The shapes of the waves of downregulated genes are also more similar in the SSCs of both genotypes (**Fig. 2**).

**FIGURE 2.**
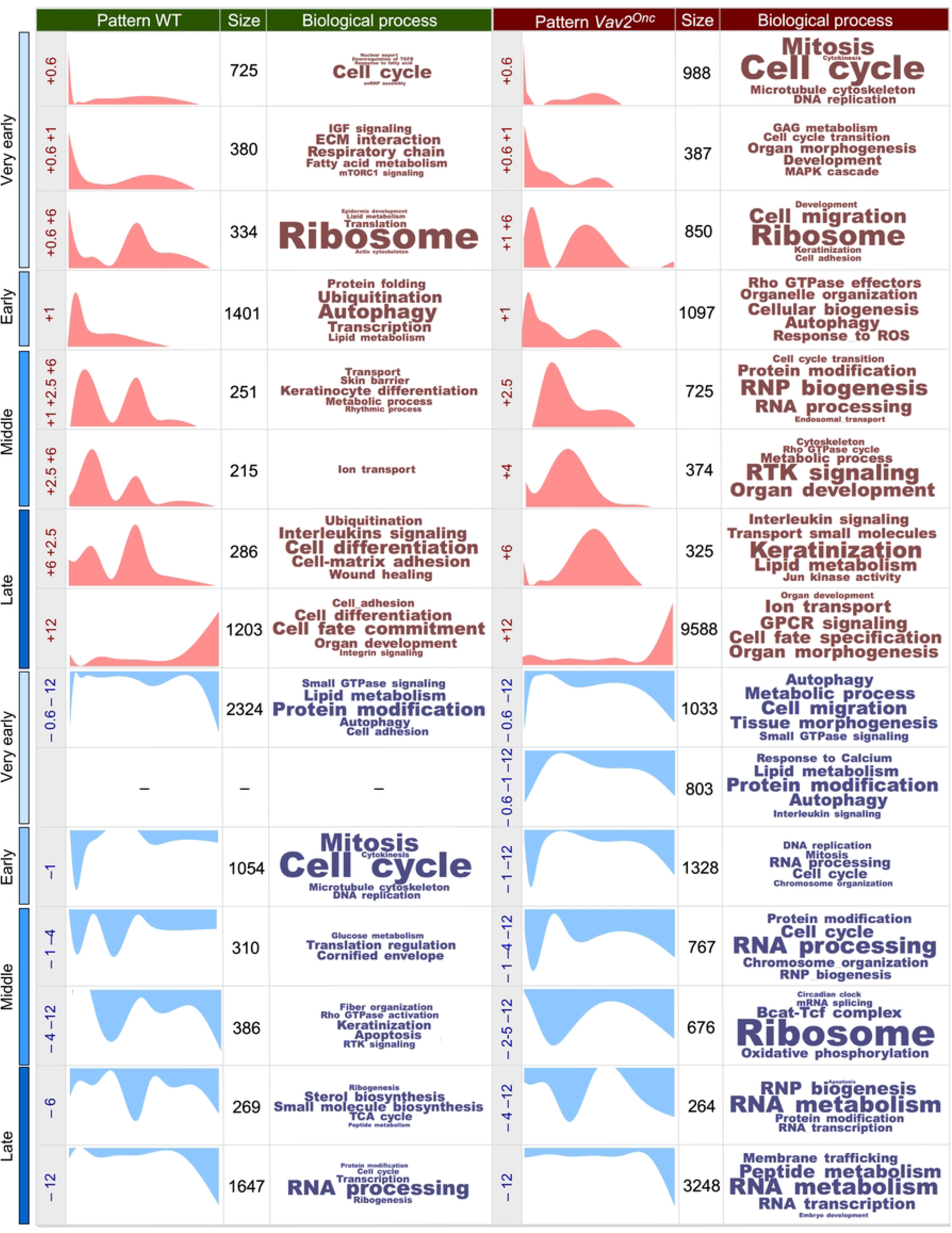
Deconvolution of the gene expression waves found in WT or *Vav2*^Onc/Onc^ SSCs. Scheme showing the expression waves found in WT SSCs (left columns, green) and *Vav2*^Onc/Onc^ SSCs (right columns, red). The following information is indicated from left to right: **(i)** the experimental time point (very early, early, middle, late) in which the indicated gene expression wave is detected; **(ii)** the time distribution of each wave, given in age of mouse in months (M), of 0.6 M (e.g., 18-days), 1 M, 2.5 M, 4 M, 6 M, and 12 M; **(iii)** a graphic representation of the indicated gene expression wave along the experimental time points used in this study; **(iv)** the number of probe sets associated with the indicated gene expression waves; and **(v)** the top-enriched biological and molecular processes (with font size proportional to the –log(*P*) obtained for each). Each wave is designated as + or – to indicate upregulated (light red waves) or downregulated (light blue waves) genes, respectively. A reference number indicating the time point at which the maximal expression peak of the wave occurs (e.g., [+6] or [–12]) is given, whereby numbers for multi-peak waves indicate the highest fold-change peak in the first position (e.g., [+1+2.5+6] shows that the highest peak was at 1 M).

We also observed significant differences in the functions associated with the gene expression waves in the SSCs of each genotype. Thus, although some functional overlap is seen at specific time points (e.g., the cell growth, ribosomal biosynthesis, autophagy-related functions, and inflammatory-related functions in waves [+0.6], [+0.6+6], [+12], and [+6+2.5]) (**Fig. 2**), the spectrum of wave-associated functions is highly genotype-dependent (**Fig. 2**). In a few cases, the same functions are detected at different time windows (e.g., ion transport is in wave [+2.5+6] in WT SSCs but in wave [+12] in *Vav2*^Onc/Onc^ SSCs; keratinocyte differentiation is in wave [+1+2.5+6] in WT SSCs but in wave [+6] in *Vav2*^Onc/Onc^ SSCs). Collectively, these results indicate that: **(i)** Vav2^Onc^ promotes a significant rewiring of the age-dependent transcriptional program of SSCs in terms of total gene numbers, time distribution of the waves of differentially expressed genes, and overall functions targeted; and **(ii)** even though Vav2^Onc^ is chronically activated in a cell stimulus-independent manner, it does not engage a fixed transcriptional program in SSCs. This suggests that, in addition to the intrinsic catalytic activity, the impact of Vav2^Onc^ on SSC biology is also influenced by other time-dependent intrinsic or extrinsic factors that contribute to shaping its impact on the transcriptome of SSCs.

Analyzing the functions associated with the gene expression waves also shed further light on the impact of Vav2^Onc^ on SSC biology. Thus, during the very early time point, the upregulated gene expression waves of WT SSCs encode functions related to cell growth, IGF (insulin growth factor) and mTORC signaling, extracellular matrix interaction regulators, and several metabolic routes. This spectrum of functions is probably associated with the expansion in the numbers of SSCs that is usually observed between postnatal days 18 and 30 in mice (18). In contrast, the waves associated with downregulation events contain genes encoding proteins involved in small GTPase signaling, cell adhesion, protein modification, and autophagy. This program is substituted as the animals age by the upregulation of genes involved in catabolic (e.g., autophagy, ubiquitination), cell differentiation, cell adhesion, and inflammatory processes (**Fig. 2**). In parallel, we observed downmodulation of genes linked to cell cycle regulation, receptor tyrosine kinase (RTK) signaling, ribogenesis, and metabolic programs (TCA cycle, sterol, and small molecule biosynthesis) (**Fig. 2**). This pattern can be linked to the post-mitotic telogen phase of the skin (which takes place in approx. 2.5-month-old mice) and with the subsequent acquisition of aging features by SSCs at later stages of the time-course analyzed in this study (7, 18). A comprehensive description of the time-dependent evolution of all these functions can be found elsewhere (18). In the case of Vav2^Onc^ SSCs, we find similar, significantly amplified functional features related to cell growth upregulation at the very early time point (**Fig. 2**). In contrast, the downregulation of cell growth-related genes that takes place at the early time point is less acute in these cells than in the controls (**Fig. 2**). At later time points, the upregulated waves of Vav2^Onc^-expressing SSCs include a large variety of functions not seen in the WT counterparts, such as signaling pathways for MAPK [+0.6+1], RTK [+2.5+6], Rho GTPase [+2.5+6], Jun-N-terminal (JNK) [+6+2.5], G-coupled protein receptors (GPCR) [+12], and ribonucleoprotein (RNP) biogenesis [+1+2.5+6] (**Fig. 2**). This indicates that, from a signaling, metabolic, and developmental perspective, the *Vav2*^Onc/Onc^ SSCs fluctuate between functional states during the time-courses analyzed, and that these fluctuations are quite different from those found in WT SSCs except at the very early time point. The functions associated with the waves of gene downregulation are also quite different between WT and *Vav2*^Onc/Onc^ SSCs (**Fig. 2**). The most conspicuous functional change is the downmodulation of RNA- and ribosomal-related functions in *Vav2*^Onc/Onc^ SSCs starting at the middle time points (**Fig. 2**). Thus, these data indicate that Vav2^Onc^ induces a large rewiring of the gene expression waves at the level of total gene numbers, time-dependent distribution of the waves, and functions involved. They also suggest that *Vav2*^Onc^ favors the stepwise engagement of different SSC functional states that are not usually seen in WT SSCs at most time points.

### Vav2^Onc^ reshapes the time-dependent expression waves present in skin stem cells

Based on the extensive transcriptomal rewiring observed in the foregoing experiments, we decided to build a network map to better visualize the changes in wave localization of the age-regulated genes in the case of WT and *Vav2*^Onc/Onc^ SSCs. In this map, the gene expression waves found in WT and *Vav2*^Onc/Onc^ SSCs are depicted as green and red circles, respectively (**Fig. 3**). In both cases, the size and darkness of the color of each wave are proportional to the total number of dynamically regulated probe sets of each wave and to the level of changes in gene probe localization seen between the WT and *Vav2*^Onc/Onc^ conditions, respectively (**Fig. 3**). The change in position of gene probes is further highlighted by the thickness of the arrows connecting the indicated WT and *Vav2*^Onc/Onc^ SSC waves (**Fig. 3**). Finally, the circles also include information on the percentage of genes that are shared with waves of the opposite genotype (white peripheral areas) or of those that are just seen in the waves of a single genotype (gray peripheral areas) (**Fig. 3**). Importantly, we also include the reference numbers assigned to the waves in **Figure 2** to facilitate their identification in **Figure 3**.

**FIGURE 3.**
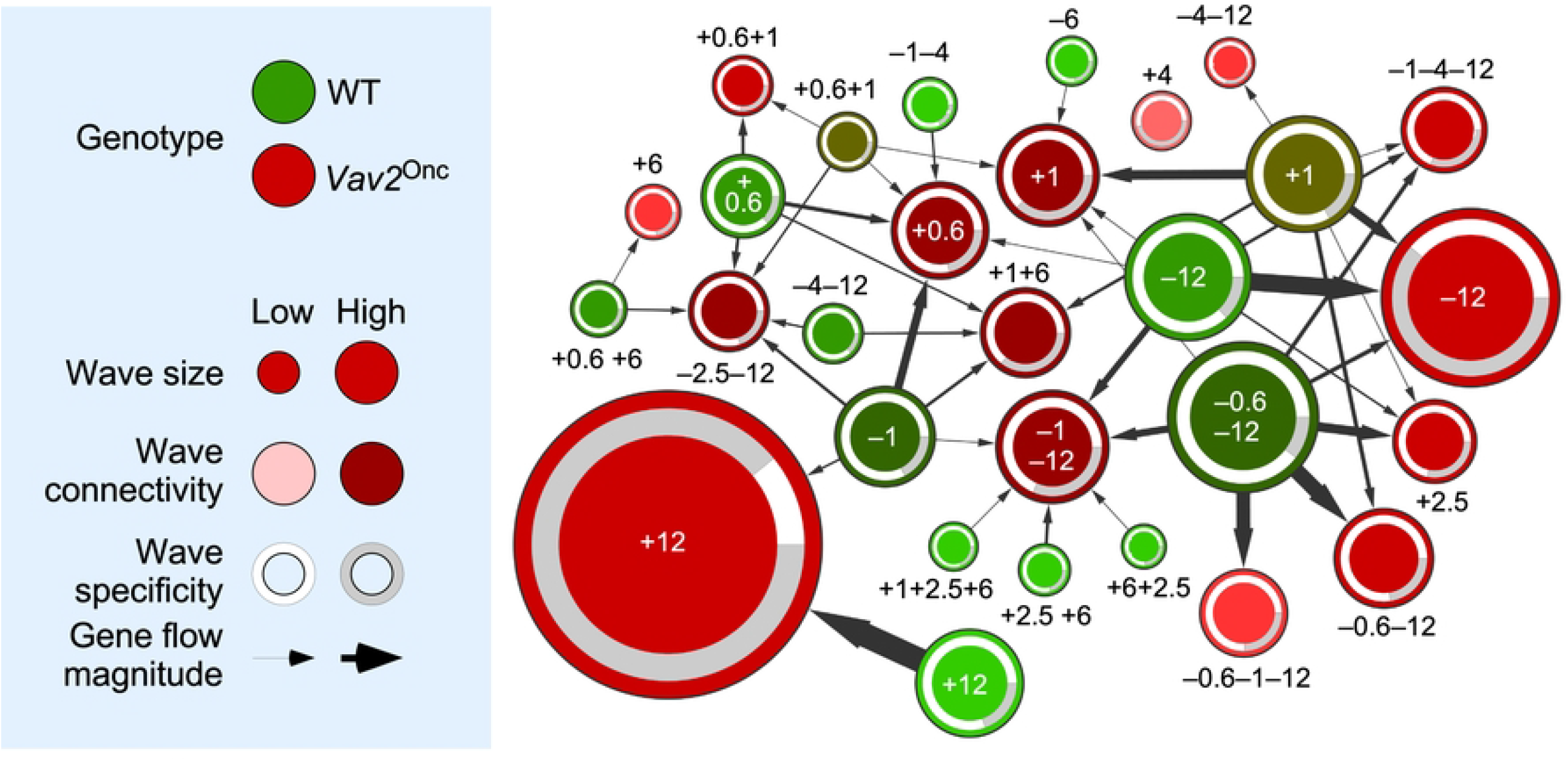
Vav2^Onc^ reshapes the gene expression waves found in SSCs. *Left*, summary of the parameters used in the network shown on the right. *Right*, dynamic flow network showing the relationship between WT (green) and *Vav2*^Onc/Onc^ (red) gene expression waves (nodes). Each expression wave is identified by the position (0.6-to-12 months old) and orientation (upregulated [+] or downregulated [–]) of its main peak(s) indicated in **Figure 2**. Wave size (number of probe sets), connectivity (number of interacting clusters in terms of probe set sharing), specificity (number of probe sets that are not dynamic in the other genotype), and flow magnitude (number of probe sets that are shared between two clusters) are indicated.

These analyses revealed extensive repositioning of transcripts between the WT and *Vav2*^Onc/Onc^ SSC waves. This phenomenon is quite significant for the *Vav2*^Onc/Onc^ waves at: **(i)** [+0.6], which also contains transcripts present in the WT waves [–1], [+0.6], [+0.6+1], [–1–4] and [+1]; **(ii)** [+1], with transcripts present in the WT waves [+1], [+0.6+1], [–6], and [– 12]; **(iii)** [+1+6], with transcripts present in the WT waves [+0.6], [–1], [–1–4], [–12], and [– 1–12]; and **(iv)** [–1–12], with transcripts present in the WT waves [+1+2.5+6], [+2.5+6], [+6+2.5], [–0.6–12], [–1], and [–12] (**Fig. 3**). The *Vav2*^Onc/Onc^ waves that were less affected with this reshuffling in the position of dynamically regulated genes include those at: **(i)** [+6], [–0.6–1–12], and [–4–12], whereby changes in position affected just one wave of each genotype; and **(ii)** [+0.6+1] and [–0.6–12], whereby changes in position involved two waves (**Fig. 3**). The *Vav2*^Onc/Onc^ wave [+4] is the only one that does not exhibit any gene originally with respect to the WT waves (**Fig. 3**). This wave is mainly composed of genes associated with RTK signaling, Rho GTPase signaling and functions, organ development, and metabolism (**Fig. 2**). Conversely, the WT waves [–0.6–12] and [+1] contained more genes that changed position in the *Vav2*^Onc/Onc^ waves (with seven and six inter-wave changes in localization, respectively) (**Fig. 3**). On the other hand, the WT waves with lower levels of gene changes include [+1+2.5+6], [+2.5+6], [+6+2.5], [–6], [–1–4], and [–12] (with a single inter-wave change in localization for each), although some content changes involved substantial numbers of transcripts (e.g., WT wave [+12]) (**Fig. 3**). These changes in wave localization are not unidirectional, as subsets of mRNAs belonging to a specific WT wave can be found in different *Vav2*^Onc/Onc^ SSC waves (**Fig. 3**). For example, the transcripts present in the WT wave [–1] are found in five different *Vav2*^Onc/Onc^ waves ([+0.6], [+1+6], [+12], [– 2.5–12], and [–1–12]) (**Fig. 3**).

Importantly, the genes involved in this transcriptional rewiring might not necessarily keep the same expression pattern in the two genotypes (e.g., genes in the upregulated WT waves [+1+2.6+6], [+2.5+6] and [+6+2.5] are found in the downregulated *Vav2*^Onc/Onc^ SSC wave [–1–12], while genes in the downregulated WT SSC wave [–6] are found in the upregulated *Vav2*^Onc/Onc^ wave [+1]) (**Fig. 3**). In other cases, different gene subsets of the same WT wave display the same and the opposite regulation in the *Vav2*^Onc/Onc^ waves (e.g., genes in the WT wave [+1] that become located in *Vav2*^Onc/Onc^ waves [+1], [+2.5], [+1+6], [–0.6–12], [–4–12], [–1–4–12] and [–12]) (**Fig. 3**). We have also found that this rewiring is highly dependent on the time-window involved. Thus, most of the *Vav2*^Onc/Onc^ waves that are distributed between the very early and middle time points contain genes that are also dynamically regulated under normal physiological conditions in WT SSCs (**Fig. 3**, see white areas in the peripheral band of each wave). In contrast, at the late time point, the *Vav2*^Onc/Onc^ waves become enriched in genes that are not found dynamically regulated in the WT waves (**Fig. 3**, see grey areas in the peripheral band of each wave). In fact, we estimated that ≈60% of all the dynamically regulated genes found in *Vav2*^Onc/Onc^ SSCs are non-dynamic in the case of WT SSCs at this time point (e.g., see *Vav2*^Onc/Onc^ SSC waves [+12] and [–12]) (**Fig. 3**). These data suggest that Vav2^Onc^ first promotes an extensive rewiring of the transcriptional program of SSCs and, subsequently, the emergence of transcriptional programs that are mostly Vav2^Onc^-dependent. This is associated with a total change in type and/or temporal distribution of biological processes in Vav2^Onc^-expressing SSCs.

### Vav2^Onc^ promotes the synchronization of pathway-associated gene signatures

We have previously shown that the time-dependent evolution of the transcriptome of WT SSCs is associated with specific patterns of co-regulation of genes signatures that are linked to well-defined hallmarks and SSC-related biological and signaling processes (18). This led us to: **(i)** investigate the impact of chronic Vav2^Onc^ signaling on the regulatory pattern of such gene signatures; and **(ii)** focus on more specific functions rather than on those usually associated with the very generic GO terms used to classify the expression waves in **Figure 2**. To this end, we utilized the transcriptomal data obtained from WT and *Vav2*^Onc/Onc^ SSCs to carry out two concatenated *in silico* steps. First, we performed co-expression analyses to identify patterns of co-regulation of the gene elements of each interrogated gene signature throughout all time points used in our study (27, 28) (**Fig. 4A**, top). In these analyses, we can expect different types of distributions based on the obtained correlation coefficients (*r*): Type 1, distributions with *r* centered around r ∼ 0, which indicates that the elements a gene signature do not follow stable coexpression trends; type 2 and 3, distributions skewed towards *r* > 0 (type 2) or *r* < 0 (type 3), indicating that a significant fraction of the genes is transcriptionally coregulated in one direction (positive or negative correlation); and, Type 4, bimodal distributions with *r* values both above and below 0, which indicate that subsets of genes of the same signature follow opposite expression patterns. The co-regulation of all gene elements of a given signature can be plotted using co-expression matrices, such as those shown in **Figure 4B**. Finally, we used single-sample gene set enrichment analyses (ssGSEA) for all the interrogated signatures and built co-expression matrices to pick up clusters of co-regulated or anti-regulated gene signatures in the transcriptome of WT and *Vav2*^Onc/Onc^ SSCs (**Fig. 4A**, bottom).

**FIGURE 4.**
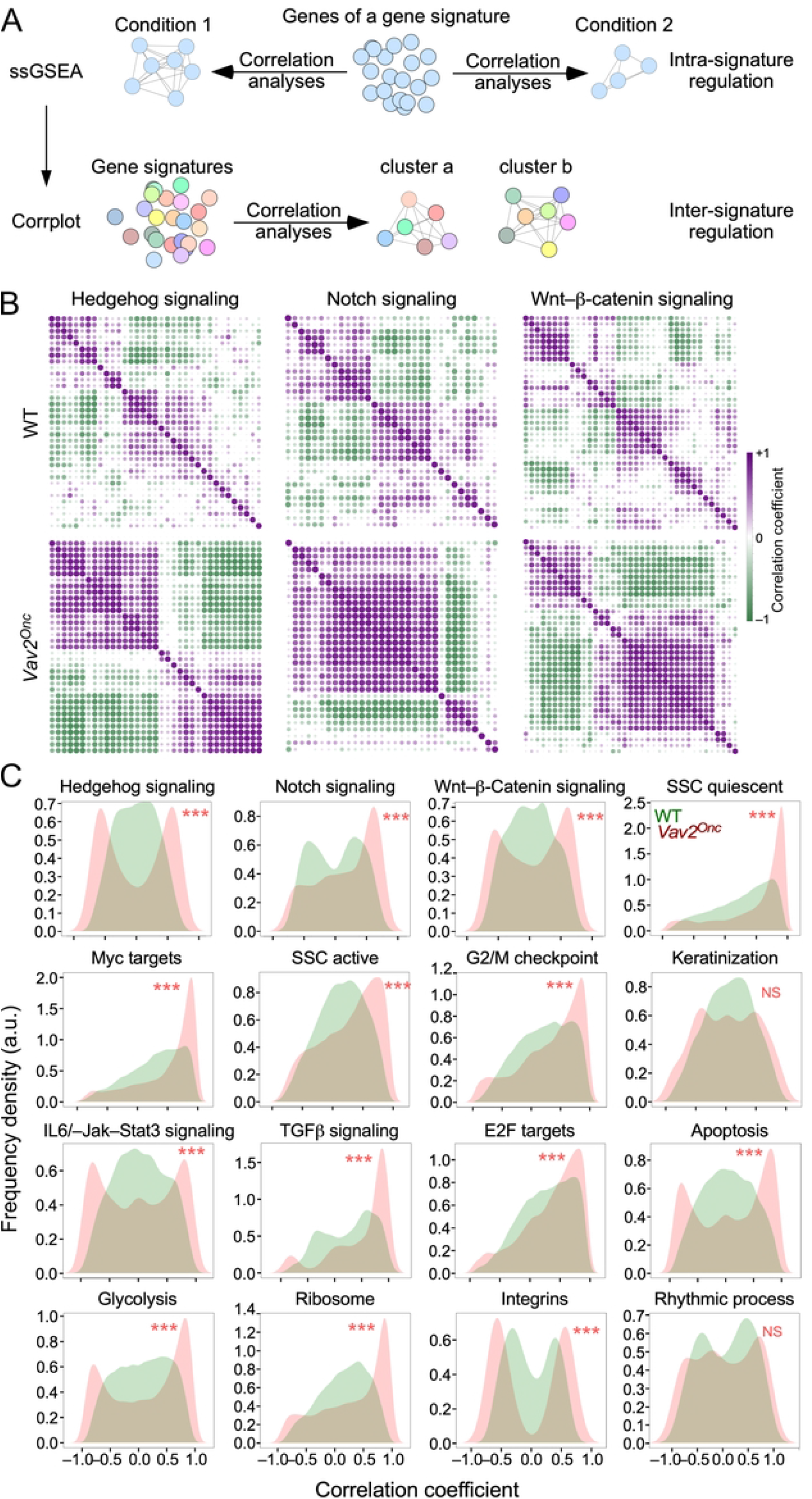
Vav2^Onc^ promotes the synchronization of pathway-associated gene signatures. **(A)** Scheme of the *in silico* analyses performed to identify intra-signature (top) and inter-signature (bottom) coregulatory events in the transcriptome of SSCs. See main text for further details. **(B)** Co-expression matrices of the transcripts integrating the indicated gene signatures in WT (top) and *Vav2*^Onc/Onc^ (bottom) SSCs. The *r* value of a given pair of gene signature elements is shown as a purple or a green dot if the two genes are co-regulated (*r* >0) or anti-regulated (*r* < 0), respectively. Dot size and color gradient reflects the magnitude of the Pearson correlation coefficient (*r*). **(C)** Density plots showing the distribution of *r* values in the coexpression matrices of indicated gene signatures in WT (green) and *Vav2*^Onc/Onc^ (red) SSCs. ***, *P* < 0.001 (Kolmogorov-Smirnov test). NS, not significant.

In the first step, we found that that the elements (single genes) of most interrogated gene signatures follow a type 1 distribution (*r* ≈ 0) (**Fig. 4B**, upper panel; **Fig. 4C**, green graphs) and, to a lesser extent, type 4 bimodal patterns (**Fig. 4C**, green graphs). Examples of each of those distributions include the hedgehog and integrin gene signatures, respectively (**Fig. 4C**, green graphs). In contrast, for *Vav2*^Onc/Onc^ SSCs, we observed a massive increase in the overall degree of co-regulation and/or anti-regulation of the elements of most gene signatures (**Fig. 4B**, lower panels). This is due to the enhancement of either the co-regulation or anti-regulation previously seen in the time-dependent transcriptome of WT SSCs (**Fig. 4B**, compare the upper and lower levels). Due to this, the co-expression patterns found in *Vav2*^Onc/Onc^ SSCs show in most cases a stronger polarization towards the type 2 (r > 0) and type 4 (r ≈ ±1) distributions (**Fig. 4C**, red graphs). Examples of each of those distributions are the elements of the epidermal stem cell quiescent signature and the hedgehog signaling signature, respectively (**Fig. 4C**, red graphs). This is a general phenomenon, as it is observed in 14 of the 16 gene signatures interrogated in these analyses (**Fig. 4C**, red graphs). The only exceptions to this trend are the gene signatures linked to keratinization and rhythmic processes (**Fig. 4C**, red graphs). These data indicate that the expression of Vav2^Onc^ increases the synchronization of gene expression of most functional gene signatures interrogated in this study.

Strikingly, using ssGSEA (**Fig. 4A**, bottom panel), we observed that Vav2^Onc^ also favors a very strong synchronization of the expression of several independent gene signatures (**Fig. 5A,B**). Consistent with this, the 10 inter-gene signature clusters previously described in WT SSCs (18) (**Fig. 5A,B**; left panels) become consolidated into just 2 clusters and a large macrocluster in *Vav*2^Onc/Onc^ SSCs (**Fig. 5A,B**; right panels). As in our previous analyses (**Fig. 3**), we observed an extensive reshuffling of the interrogated gene signatures from specific clusters of WT SSCs to the new clusters found in *Vav2*^Onc/Onc^ SSCs (**Fig. 5A**, see localization of indicated gene signatures between the left and right correlation matrices). In fact, there is only one cluster of co-regulated gene signatures that is identical in the SSCs of both genotypes (namely, clusters c_WT_ and b_Onc_; **Fig. 5**). This cluster harbors co-regulated gene signatures associated with epidermal development, epidermal cell differentiation, and keratinization (**Fig. 5B**). In addition to this extensive reshuffling, we have also found very limited cases of either loss or gain of new co-regulated gene signatures in the clusters found in *Vav*2^Onc/Onc^ SSCs. Those include the loss of the Wnt–β-catenin and TNFα–NFκB signaling gene signatures (**Fig. 5B**, left column, red font) and the *de novo* incorporation of a protein translation-associated gene signature to the cluster designated as c_Onc_ in *Vav*2^Onc/Onc^ SSCs (**Fig. 5B**, right column, red font).

**FIGURE 5.**
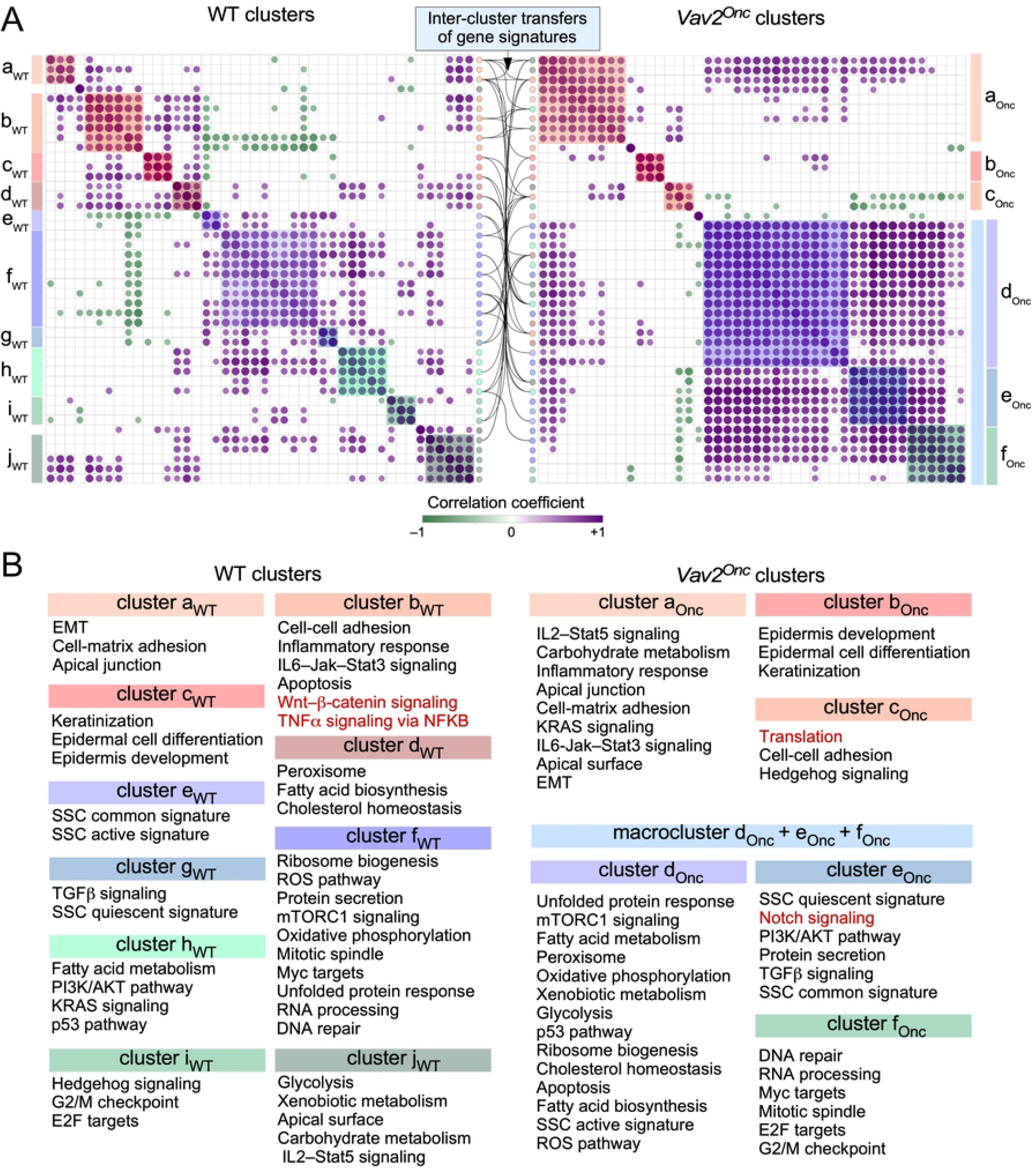
Vav2^Onc^ promotes synchronization of multiple gene signatures in SSCs. **(A)** Co-expression matrices showing the correlation across time among the enrichment scores of the gene expression signatures shown in (B) in WT SSCs (left) and *Vav2*^Onc/Onc^ SSCs (right). Only correlations with *P ≤* 0.05 have been considered as statistically significant. Positive and negative correlations are shown in purple and green, respectively. Dot size and color gradient reflects the magnitude of the r value obtained for each indicated pair. Pathway clusters are indicated with color shades. The position of a given gene expression signature in the two matrices are traced lines between the two co-expression matrices (inter-cluster transfers of gene signatures). The co-expression matrix for the WT SSC gene signatures (A, left panel) is taken from [18]. **(B)** Gene signatures co-regulated in the same clusters in WT SSCs (left columns) or *Vav2*^Onc/Onc^ SSCs (right column). Red indicates gene signatures that are not conserved in the clusters of the opposite genotype SSCs.

### Vav2^Onc^ rewires the transcription factor landscape in skin stem cells

We assumed that the extensive age-dependent transcriptional rewiring observed in *Vav2*^Onc/Onc^ SSCs had to be associated with changes in the normal transcription factor landscape of skin stem cells. To investigate this, we next performed weighted correlation network analyses (WGCNA) to calculate hierarchical co-expression relationships among all the genes identified in the expression waves found in both WT and mutant SSCs (**Fig. 2**). This approach entailed the calculation of the adjacency and the intramodular connectivity for all the wave gene components (**Fig. 6A**, top panels), two parameters that are highly related to the Eigengene-based cluster membership score of each transcript. To further increase the stringency of these analyses, we defined hubs as only the 10% of the top-connected nodes in each age period-specific gene expression wave (**Fig. 6A**, bottom graphs). We found that only 26.5% of the transcriptomal hubs are conserved between WT and *Vav2*^Onc/Onc^ SSCs, which is consistent with the extensive transcriptomal rewiring observed in **Figures 2** to **4**. This reduced overlap is seen at the very early (23.9% of shared hub elements), early (17.7% of shared hub elements), middle (29.7% of shared hub elements), and late (6.65% of shared hub elements) experimental time points (**Fig. 6B**, blue colors). The very low level of overlap in the hub elements between WT and *Vav2*^Onc/Onc^ SSCs seen in the late time-window is consistent with the increase of Vav2^Onc^-specific transcriptional programs previously observed in our *in silico* analyses (**Figs. 2** and **3**). These findings indicate that the overall transcriptional rewiring induced by Vav2^Onc^ in SSCs is accompanied by a parallel reshaping of the overall transcriptomal hub landscape.

**FIGURE 6.**
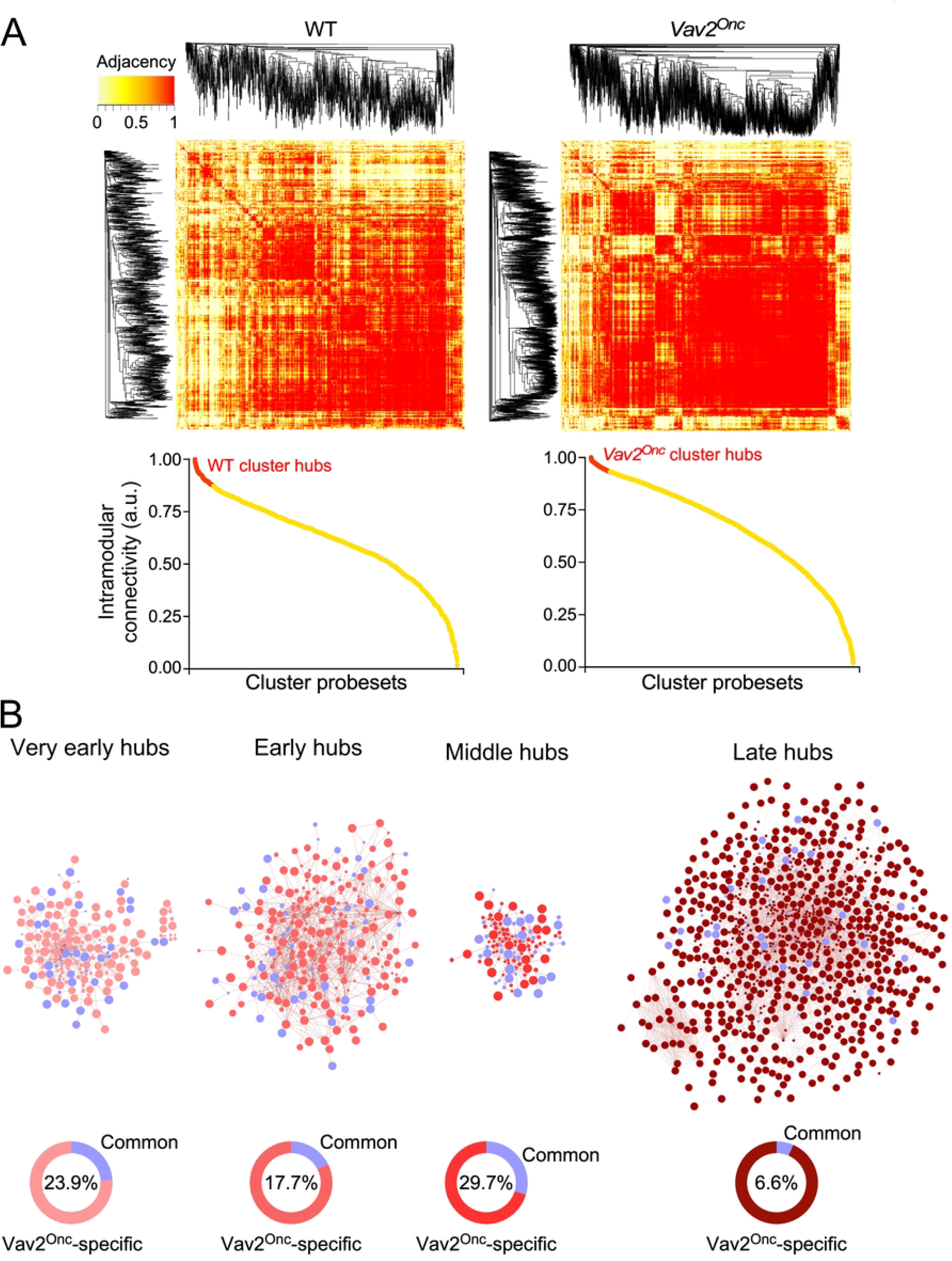
Vav2^Onc^ promotes new transcriptomal hubs in SSCs. **(A)** *Top*, representative examples of the adjacency heatmaps computed for each expression pattern applying the WGCNA algorithm to the WT SSC or *Vav2*^Onc/Onc^ SSC transcriptome. Red areas indicate the presence of probe sets that are highly connected transcriptionally. Probe set clustering according to adjacency is depicted on the sides of the heatmap. *Bottom*, representative examples of dot plots showing the intramodular connectivity score for each of the probe sets belonging the coexpression clusters analyzed in the top panel. Transcriptional hubs (highlighted in red) are defined as the 10% most interconnected probe sets. Please, note that the diagram for the WT SSC hubs has been used before (18). **(B)** *Top*, transcriptomal hubs found in *Vav2*^Onc/Onc^ SSCs at the indicated experimental time points. The hubs shared with WT are shown in blue color. *Bottom*, percentage of hubs that are specific to *Vav2*^Onc/Onc^ SSCs (red) or common to both *Vav2*^Onc/Onc^ and WT SSCs (blue) (blue). The percentage of hubs shared by SSCs of both genotypes is given inside each graph.

We hypothesized that the transcription factors responsible for the Vav2^Onc^-mediated transcriptomal rewiring had to be contained within the hubs described above. In line with this idea, we found that the landscape of transcription factors acting as hubs was very different between WT and *Vav2*^Onc/Onc^ SSCs (**Fig. 7A**). In WT SSCs, they include members of the zinc finger C2H2, HOX-like, and E2F families (**Fig. 7B, Table S2**). In contrast, in *Vav2*^Onc/Onc^ SSCs, they encompass specific members of the basic helix-loop-helix, Forkhead box, Gata, nuclear hormone receptor, Tcf, Ets, Nkl, Hox-like, Stat, and pluripotency-associated families (**Fig. 7B, Table S2**). We reasoned that if these transcription factors mediate the Vav2^Onc^-induced transcriptomic remodeling, they had to target many of genes present in the Vav2^Onc^-driven expression waves. Consistent with this idea, we found that most of the dynamically regulated genes in *Vav2*^Onc/Onc^ SSCs (**Fig. 2**) harbor site(s) for these transcription factors (**Fig. 7B,C**; **Table S2**). This is Vav2^Onc^-specific, since these sites are not enriched in the case of the dynamically-regulated genes in WT SSCs (**Fig. 7B, C**; **Table S2**). Indeed, our microarray data indicate that Vav2^Onc^ transcriptionally regulates the activity of a large fraction (55%) of these transcription factors (**Fig. 7D**, left panel). Importantly, ssGSEA analyses indicate that these Vav2^Onc^-driven transcription factors show high expression levels in quiescent SSCs and become strongly downmodulated upon SCC activation and differentiation (**Fig. 7E**, top panel). As a control, performing these analyses using equally sized random collections of transcription factors shows no significant enrichments (**Fig. 7E**, bottom panel).

**FIGURE 7.**
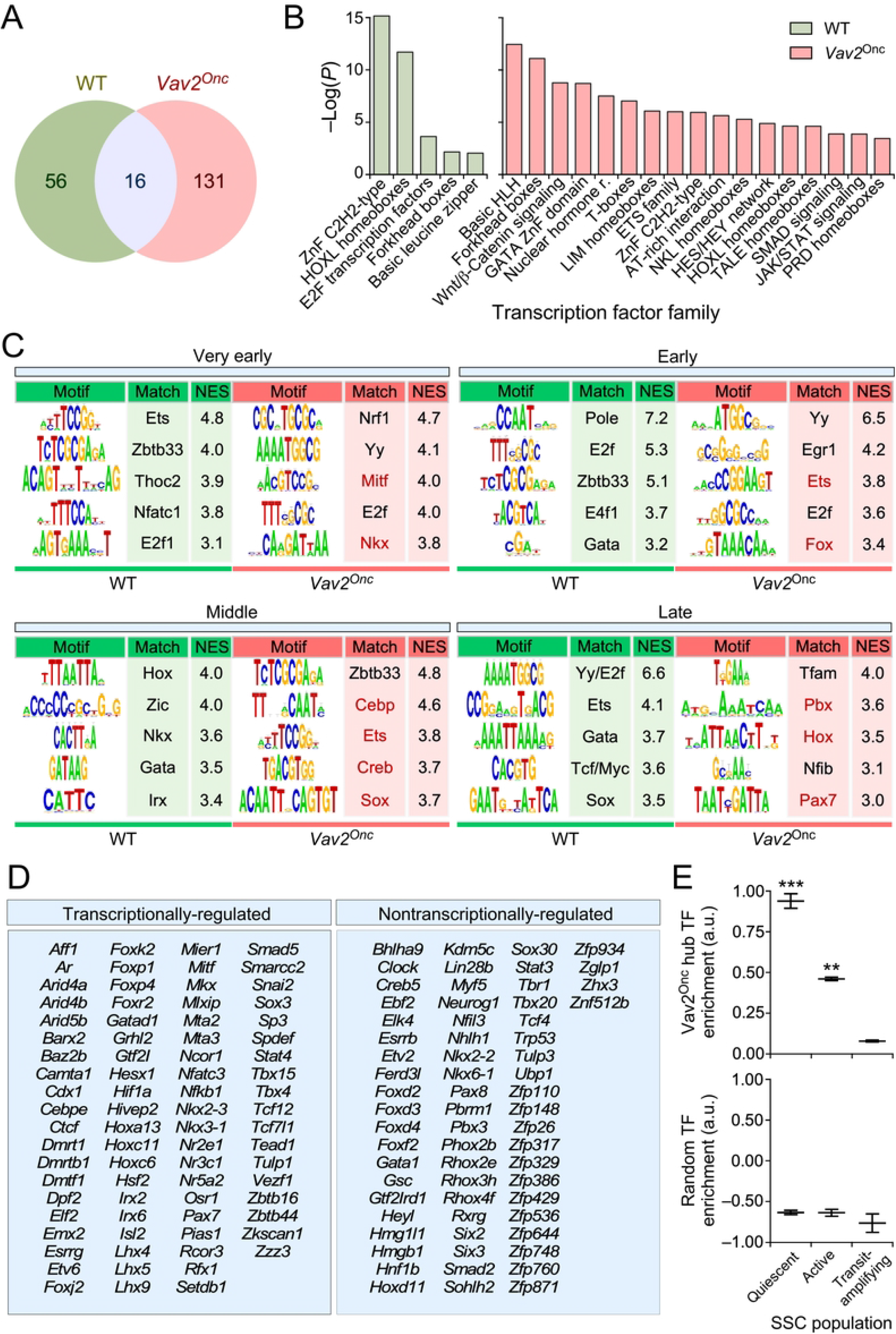
Vav2^Onc^ rewires the transcription factor (TF) landscape in SSCs. **(A)** Venn diagram showing the overlap in TF identify in the transcriptomal hubs of WT or *Vav2*^Onc/Onc^ SSCs. **(B)** Main TF families enriched in the WT or Vav2^Onc^ collections of TF hubs. For all, *P ≤* 0.05 (Fisher’s exact test). **(C)** The main enriched TF binding sites in the promoters of time-regulated genes from WT or *Vav2*^Onc/Onc^ SSCs. Red indicates sites for which the TF is in a Vav2-regulated transcriptional hub. Normalized enrichment scores (NES) are indicated. For all, *P ≤* 0.05 (iRegulon Wilcoxon rank-sum paired test). **(D)** List of TFs found in the transcriptomal hubs of *Vav2*^Onc/Onc^ SSCs. TFs that are differentially expressed at any time point as compared to WT SSCs are labeled as “transcriptionally-regulated” (left column). **(E)** Plots showing the enrichment of the Vav2^Onc^ hub TFs (top) and of an equally sized random collection of TFs (bottom) in the indicated epidermal cell populations. **, *P* < 0.01; ***, *P* < 0.001 against TAC population (Student’s *t*-test). Data represent median, minimum, and maximum values. TAC, transit-amplifying cell.

## DISCUSSION

Given their stemness condition, it could be inferred that SSCs have to remain in a “platonic-like”, stable functional state throughout the life of an organism unless they are required for regenerative responses. However, we have recently shown that approximately one-third of the transcriptome of mouse SSCs undergoes quite significant age-dependent variations under normal tissue homeostatic conditions. This is associated with well-defined age-dependent gene expression and functional waves (18). Likewise, an independent study has demonstrated extensive changes in the transcriptome of adult (18-month-old) SSCs (7). It is likely that these transcriptome changes are dependent on internal SSC clocks as well as on external signals from the surrounding cells of the SSC bulge niche. In addition to the time dimension, the functional features of SSC are also highly influenced by the activation state of specific signal transduction proteins. Thus, we have shown in a recent study that interfering with the catalytic function of Vav2, using either a gain-of-function or a loss-of-function approach leads to extensive changes in the numbers, regenerative capacity, and the overall transcriptome of SSCs isolated from 2.5-month-old mice. Such changes are connected to increasing proliferation while maintaining the biological programs linked to SSC pluripotency (17). However, given that these analyses have been carried out at a single age time point (17), it is still unclear how Vav2 signaling influences the functional state of SSCs throughout the lifespan of the animals. Here, we have now approached this issue by examining the effect of constitutive Vav2^Onc^ signaling in the transcriptome of SSCs isolated from six independent age points in mice.

Our data indicate that the constitutive signaling of Vav2^Onc^ has a very profound impact on the overall transcriptome of SSCs. Thus, we observed much higher numbers of age-regulated genes in *Vav2*^Onc/Onc^ SSCs (64.6%) than in WT SSCs (31.9%) (**Fig. 1D**). Many of these genes represent *de novo* regulatory events that are not found in the age-dependent transcriptome of WT SSCs (**Fig. 1E**). Perhaps more importantly, we have found a few interesting features of this Vav2^Onc^-regulated transcriptome. On the one hand, and despite the constitutively catalytic activity exhibited by Vav2^Onc^, we have found that the Vav2^Onc^-regulated transcriptome is still organized in age-dependent waves of gene expression rather than being fixed over time (**Figs. 1F** and **2**). On the other hand, we observed that Vav2^onc^ elicits an extensive rewiring of the gene expression waves that are normally observed in SSCs. This rewiring involves: **(i)** the age-dependent expression of genes not found in SSCs (**Figs. 2** and **3**); **(ii)** extensive changes in the time-dependent wave position of genes as compared to WT SSCs (**Fig. 3**); **(iii)** in some cases, changes in the expression pattern of the dynamically regulated genes (e.g., changing from up- to downregulation) (**Fig. 3**); **(iv)** an increased level of co- or anti-regulation of genes belonging to gene signatures associated with hallmark biological processes and SSC-related states (**Fig. 4**); and **(v)** a further synchronization in the co- or anti-regulation of independent functional gene signatures in few co-regulated clusters (**Fig. 5**). As a result of this extensive rewiring, very different age-dependent functional states can be recognized in the Vav2^Onc/Onc^ SSCs as compared to the WT counterparts (**Fig. 2**). In general, these states are associated with larger proliferative responses in the very early postnatal phase and more active and enriched signaling/metabolic programs in the subsequent time points analyzed. It is expected therefore that, as is the case of 2.5-month-old mice (17), the SSCs from older *Vav2*^Onc/Onc^ mice would have an increased regenerative potential as compared to that of control mice.

It is difficult to decipher the master transcriptional regulators that underlie this extensive transcriptomal rewiring using wet-lab approaches. To shed light on this issue, we used *in silico* approaches based on the identification of transcriptional hubs in the Vav2^Onc^-regulated gene expression program and, subsequently, the search for binding sites for those transcriptional hubs in the promoter regions of Vav2^Onc^-regulated genes. Using this approach, we have found that Vav2^Onc^ promotes an extensive rewiring and amplification of the age-dependent transcription factor landscape present in SSCs (**Fig. 7**). This landscape includes transcription factors of the Cebp, Creb, Ets, Fox, Hox, and Sox families, many of which have been previously linked to the regulation of SSC identity and homeostasis (28, 29). Interestingly, these Vav2^Onc^-engaged transcriptional hubs are specifically enriched in quiescent SSCs and, to a lesser extent, active SSCs, but not in subsequent epidermal cell differentiation stages. In addition to transcriptomal hubs whose expression is specifically regulated in Vav2^Onc/Onc^ SSCs, we have found other hubs that do not undergo statistically significant changes in expression between *Vav2*^Onc/Onc^ and WT SSCs (**Fig. 7D**). This suggests that a significant part of the Vav2^Onc^-regulated transcription factor landscape could be regulated by posttranscriptional mechanisms. Future studies using wet-lab approaches will be needed to further validate this potential Vav2^Onc^-regulated transcription factor landscape and pinpoint the most critical elements from a regulatory point of view.

Data from our previous study (17) and the present work suggest that Vav2^Onc^ will promote increased numbers of SSCs endowed with more basal regenerative potential for longer periods of time in mice. However, they also give a warning sign, since it is foreseeable that this potentially beneficial feature could represent a problem when oncogenic mutations arise in those cells. In fact, we have previously demonstrated that Vav2^Onc^ promotes the generation of a protumorigenic, hyperplastic niche in the skin that can facilitate tumor formation upon the emergence of genetic lesions in the keratinocyte cell layer of the skin (19). On the brighter side, it is possible to envision that the Vav2^Onc^-associated transcriptome of SSCs will have specific Achilles’ heels that can be targeted to hamper the fitness of cancer SSCs. Such an avenue has been, in fact, demonstrated in the case of more differentiated epithelial components of the skin (19, 30).

## MATERIALS AND METHODS

### Ethics statement

All mouse experiments were performed according to protocols approved by the Bioethics Committee of the University of Salamanca and the animal experimentation authorities of the autonomous government of Castilla y León (Spain). No patients or patient-derived samples were used in this work.

### Animal studies

Control and *Vav2*^Onc/Onc^ mice (19, 20) had a C57BL/6J genetic background. Animals were kept in ventilated rooms in a pathogen-free facility under controlled temperature (23 °C), humidity (50%) and illumination (12-hour-light / 12-hour-dark cycle) conditions.

### Skin stem cell isolation

Epidermal stem cells were isolated as previously described (31). Briefly, mouse back skin was digested in 0.25% trypsin (Thermo Fisher Scientific, cat. # 25200056) overnight at 4 °C to collect the keratinocytes from the epidermis. This cell suspension was then filtered, resuspended in EMEM (Lonza, cat. # BE06-174G) supplemented with 15% fetal bovine serum (Thermo Fisher Scientific, cat. # 10500064) and incubated for 30 min on ice with biotin-conjugated antibodies to CD34 (1:50, eBioscience, cat. # 13-0341-85) followed by another 30 min incubation with APC-conjugated streptavidin (1:300, BD Biosciences, cat. # 554067) and PE-conjugated antibodies to CD49f (1:200, AbD Serotec, cat. # MCA699PE). Cells were stained with DAPI (4’,6-diamidino-2-phenylindole) (0.1 ng/μL, Sigma-Aldrich, cat. # D9542) for 5 min to exclude dead cells. Cells positive for CD34 and CD49f were isolated using a FACSAria III flow cytometer (BD Biosciences) and analyzed with the FlowJo software.

### RNA extraction and transcriptome profiling

SSCs were lysed in RLT buffer (QIAGEN, cat. # 74004), and RNA was extracted using the QIAGEN RNeasy Micro Kit (QIAGEN, cat. # 74004) according to the manufacturer’s instructions. Purified RNA was processed as indicated elsewhere (32) and hybridized to Affymetrix GeneChip Mouse Gene 1.0 ST microarrays. R (version 4.1.2). The Perl software (version 5.26.2) was used to perform the bioinformatic analyses as previously described (18). Signal intensity values were obtained from CEL files after applying the Robust Multichip Average (RMA) function from the ‘affy’ package for background adjustment, quantile normalization and summarization (33).

### Establishment of gene expression patterns

Both Chi–squared and fold-change (FC) thresholds were used to distinguish probe sets with dynamic and stable behavior along the time–points interrogated, as previously reported [16]. Briefly, for each probe set, a chi–squared test with N–1 degrees of freedom were applied as follows:

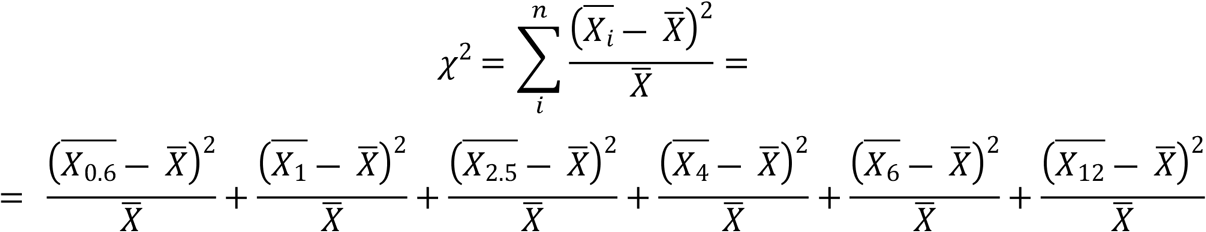

where 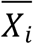 is the mean expression of the gene for the triplicates for each time point *i*, 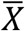 is the overall mean expression across all time points, and *n* is the number of time points. For probe sets satisfying *P* (*χ*^2^) < 0.01, a FC ≥ 2 requirement was empirically established. Unsupervised soft clustering was performed using the Mfuzz R package (34); this was used to identify time-course expression profiles. After the expression values were standarized for Euclidian space clustering, the *mfuzz* function with a fuzzifier value of 1.25 and a ranging number of cluster centers was used to determine the optimal number of non-overlapping expression patterns. For probe set inclusion in a particular cluster, the membership value threshold was set to 0.5. The resulting expression profiles were named according to the positions of the main positive (+) and negative (–) enrichments (peaks), i.e., 0.6 = 0.6-month-old; 1 = 1-month-old: 2.5 = 2.5-month-old; 4 = 4-month-old; 6 = 6-month-old; and 12 = 12-month-old. To evaluate how Vav2^Onc^ activity affects SSC transcriptional dynamics, the fluctuations of each dynamic probe set between the WT genotype (data from (18)) and the *Vav2*^*Onc/Onc*^ genotype (data from this study) were analyzed. Using Cytoscape software (35), the results were represented as a network, with nodes indicating expression patterns and edges (arrows) representing how dynamic transcripts behave in the Vav2^Onc^ context as compared to the WT. Node color, color hue, and node size were set according to pattern genotype, connectivity, and size, respectively. The inner ring indicates the dynamic probe sets within the pattern that are not dynamic in the other genotype. Edge thickness positively reflects the amount of transcripts that share the same behavior. The sensitivity for edge representation was set to 10% of the WT pattern size. For visualization of expression and enrichment data in plots, a 0–1 normalization was used as follows:

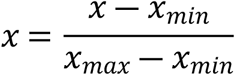

### Gene set annotation and enrichment analyses

Differentially expressed genes between genotypes were identified using linear models for microarray data (limma) (36), adjusting *P* values for multiple comparisons by applying the Benjamini–Hochberg correction method (37). Functional annotation was performed using Metascape (38) for biological processes, and ToppFun (39) for molecular functions. An FDR q-value of 0.05 was set as threshold for statistical significance. Single-sample gene set enrichment analysis (ssGSEA) (40, 41) was used to calculate the fitness of different hallmark, gene ontology, and SSC signatures across time and genotypes. Hallmark and gene ontology signatures were obtained from the Molecular Signatures Database (27). The SSC gene sets were obtained from (28). The time-course enrichment scores for these signatures were used to build the signature correlation matrix, calculated through *corrplot* (https://github.com/taiyun/corrplot). Correlations were considered statistically significant when the Pearson coefficient corresponded to *P ≤* 0.05. Functional clusters were established when every pairwise correlation within a group of signatures was found significant.

### Weighted correlation network analyses

To find the best representatives of every gene expression pattern, the *WGCNA* R package was used (42). Each weighted gene network was constructed from the corresponding expression matrix using the *blockwiseModules* function. The *pickSoftThreshold* function was used to select the soft thresholding power according to network topology. Consensus module detection within each expression pattern was omitted and kept to one module, as the number of clusters had been already optimized. The heatmap plot depicting the adjacency matrix was created with the *TOMplot* function. To calculate the intramodular connectivity for each gene, the whole network connectivity was computed for each expression pattern through the *intramodularConnectivity* function. Hubs were defined as the 10% most connected genes within each expression pattern. The known functional interactions among hubs were obtained through the String tool (43) and represented using Cytoscape (35). Transcription factor classification was performed using ToppFun (39). The iRegulon software was used to determine the transcription factor binding motifs in the promoters of the co-regulated genes (44). A collection of 9713 position weight matrices (PWMs) was applied to analyze 10 kb centered around the TSS. With a maximum false discovery rate (FDR) on motif similarity below 0.001, we performed motif detection, track discovery, motif-to-factor mapping, and target detection.

### Statistics

The type of statistical tests performed, and the statistical significance, are indicated for each panel either in the figure legends or in the main text. Data normality and equality of variances were analyzed with Shapiro-Wilk and Bartlett’s tests, respectively. Parametric distributions were analyzed using Student’s *t*-test. Sidak’s multiple comparison test was used when comparing different sets of means. The chi-squared test was used to determine the significance of the differences between expected and observed frequencies. The Kolmogorov-Smirnov test was used to compare probability distributions. In all cases, values were considered significant when *P ≤* 0.05. Data obtained are given as the mean ± SEM unless otherwise indicated.

## DATA AVAILABILITY

Microarray data corresponding to WT SSCs were obtained from the GEO database (https://www.ncbi.nlm.nih.gov/geo/) entry GSE137176 (18). The Vav2^Onc^ data reported in this paper were merged with the former and deposited under the accession number GSE140152.

## MATERIALS AVAILABILITY

All relevant data are available from the corresponding author upon reasonable request. A Materials Transfer Agreement could be required in the case of potential commercial applications.

## AUTHOR CONTRIBUTIONS

L.F.L.-M. did the experimental work and the bioinformatic analyses, performed artwork design, and wrote the manuscript. X.R.B. conceived the work and carried out the final editing of the manuscript and figures.

## ACKNOWLEDGEMENTS

We thank A. Abad for animal-based work, M. Blázquez for laboratory work, A. Prieto for FACS-related technical assistance, and E. Fermiñán for microarray-hybridization procedures. X.R.B.’s work has been supported by Worldwide Cancer Research (14-1248), the RTI2018-096481-B-100 grant cofunded by MCIN/AEI/10.13039/501100011033 and the European Research Development Fund “A way of making Europe” of the European Union, the Spanish Association against Cancer (GC16173472GARC), the Castilla-León autonomous government (CSI145P20, CLC-2017-01), and “la Caixa” Banking Foundation (HR20-00164). X.R.B.’s institution is supported by the Programas de Apoyo a Planes Estratégicos de Investigación de Estructuras de Investigación de Excelencia of the Castilla-León government (CLC-2017-01 and CL-EI-2021-02) that were both cofounded by the European Research Development Fund. L.F.L.-M. contract has been supported by funding from the Spanish Ministry of Education, Culture and Sports (FPU13/02923) and, subsequently, by the CLC-2017-01 grant.

